# Deep-Tissue Two-Photon Brain Imaging Enabled by a Tunable Fiber-Optic Dispersive Wave Generator

**DOI:** 10.1101/2025.03.20.644353

**Authors:** Marvin Edelmann, Andreu Matamoros-Angles, Mohsin Shafiq, Mikhail Pergament, Markus Glatzel, Franz X. Kärtner

**Affiliations:** Center for Free-Electron Laser Science CFEL, Deutsches Elektronen-Synchrotron DESY, Notkestr. 85, 22607 Hamburg, Germany; Department of Physics, Universität Hamburg, Jungiusstr. 9, 20355 Hamburg, Germany; Max Planck School of Photonics, Hans-Knöll-Str. 1, 07745 Jena, Germany; Institute of Neuropathology, University Medical Center Hamburg-Eppendorf (UKE), Martinistr. 52, 20246 Hamburg, Germany

## Abstract

Here, we present a fiber-optic dispersive wave generator for highly-efficient, wavelength-tunable ultrashort pulse generation, enabling multicolor deep-tissue two-photon imaging of neuronal and vascular structures in labeled mouse brain. Guided by comprehensive numerical simulations, a compact Yb:fiber laser-driven system is constructed that utilizes precisely parameter- and phase-matching-controlled dispersive wave generation in a photonic crystal fiber. The system delivers sub-100 fs pulses with over ∼6.7 nJ of energy across a continuously tunable spectral range of 880–950 nm, achieving a record-high optical conversion efficiency of up to 65%. Optimizing the output for two-photon excitation of enhanced Green Fluorescent Protein and SYTOX Orange enables high-resolution structural imaging in mouse hippocampus and cerebellum at depths exceeding 450 µm. This technique for wavelength-tunable, high-energy and ultrashort pulse generation with record optical efficiency represents a significant advancement in ultrafast fiber laser technology for versatile biomedical two-photon imaging applications.

## 1. Introduction

Two-photon microscopy (2PM) enables high-resolution, three-dimensional imaging in scattering biological tissue at large imaging depths, making it a crucial tool in modern life-sciences such as embryology [1], oncology [2], and neuroscience [3]. The functionality of 2PM thereby strongly dependents on advancements in ultrafast laser technology, in particular the development of high-performance femtosecond laser systems that spectrally match the nonlinear excitation profiles of application-specific fluorophores and biomolecular markers. In the field of neuroscience for instance, the visualization of neuronal morphology and cell dynamics via 2PM largely relies on pulsed laser excitation in the spectral window between 800 – 1000 nm [5]. Prominent examples of important neuronal labelling are the derivatives of green fluorescent protein (GFP) with two-photon excitation peak near 920 nm, utilized, e.g., in the form of EGFP-expressing transgenic mouse models to study neuronal connectivity, synaptic plasticity, and neural activity dynamics in both healthy and diseased brain states [6-9]. There are further a series of chemical and genetically encoded calcium indicators, such as Fluo-4 AM and the GFP-based GCaMP variants, with two-photon excitation peaks between 900 – 950 nm, which are vital tools for capturing neuronal activity by detecting intracellular calcium transients associated with the neurons action potentials [10-13]. In addition, there are widely applied fluorescent probes such as Alexa Fluor and Fluorescein derivatives (as example AF488 with excitation peak at 950 nm), which are also extensively used e.g., to visualize specific protein markers associated with neural and vascular structures [14-16].

Precisely matching said fluorophore and neuronal markers excitation profiles offers significant advantages, including optimized fluorescence signal for enhanced image brightness and contrast, improved imaging depth, and a reduced probability of photodamage due to lower required laser power [17,18]. Traditionally, solid-state Ti:Sapphire lasers have been standard systems in this context, offering Watt-level average power and sub-100 fs pulses over a wide spectral range of 700–1000 nm [19,20]. However, this performance comes with high costs, significant technical complexity, large and bulky physical footprints, the need for external cooling, and demanding maintenance and alignment requirements. In recent years, fiber-optic laser systems have therefore emerged as rapidly advancing, cost-efficient, compact, and low-maintenance alternatives [21,22]. Today, Neodymium and frequency-doubled Thulium-doped fiber lasers routinely generate high-power, sub-150 fs pulses between 900-920 nm [23-25]. However, due to the fundamentally limited gain bandwidth of the dopants and a necessity for highly-optimized, parameter-bound working points, their performance is typically limited to a single, fixed center wavelength e.g., 910 nm or 920 nm.

To achieve broadband wavelength-tunability with fiber-optic laser systems, a common approach is to utilize excessive nonlinear spectral broadening, e.g., in highly nonlinear photonic crystal fibers (PCF). For example, Chou et al. recently demonstrated a broadband tunable Yb:fiber (Ytterbium-doped fiber) laser source operating in the 740 – 1240 nm spectral range for 2PM applications [26], using self-phase modulation enabled spectral selection (SESS) in a PCF [27]. However, due to the limited conversion efficiency of 10-20% that is characteristic for SESS-based 2PM sources [28-30], the obtained pulse energies in the 800 – 1000 nm range are restricted to below 2 nJ, which is insufficient for deep-tissue two-photon brain imaging applications with most 2PM setups. As possible alternative, Zach et al. recently demonstrated an Er:fiber (Erbium-doped fiber) laser source that employs soliton self-frequency shift (SSFS) and subsequent frequency doubling to generate 120 fs pulses with more than 500 mW average power at 80 MHz repetition rate, tunable in the 810 – 1000 nm range [31]. However, this performance comes at a cost at a significantly increased technical complexity e.g., through the usage of spiraled large-mode area fibers with required high-order mode suppression and a complicated pumping scheme based on an external, high-power Raman-laser.

In this work, we present a nonlinear fiber-optic laser system for highly efficient, tunable femtosecond pulse generation between 880 – 950 nm that is tailored for high-performance deep-tissue neural two-photon imaging applications. The underlaying physical mechanism is based on precise parameter and phase-matching control to enable wavelength-tunable, highly efficient dispersive wave (DW) generation in a PCF. Guided by comprehensive numerical simulations, a compact, fiber-optic DW generator is constructed, able to generate >6.7 nJ, sub-100 fs pulses with up to 65% optical conversion efficiency from a Yb:fiber laser, continuously wavelength-tunable in the 880 – 950 nm spectral range. The system is subsequently applied for versatile deep-tissue two-photon brain imaging. Using the 920 nm DW output, neuronal and vascular structures in the hippocampus of a transgenic GFP-expressing mouse are visualized to a depth of 600 µm. The DW generator is further applied at 950 nm center wavelength, to enable the resolution of individual neuronal nuclei and blood vessel in the granular and molecular layers of a Sytox Orange labeled mouse cerebellum down to 450 µm imaging depth.

## 2. Physical Mechanism and Numerical Simulations

Comprehensive numerical simulations are applied to illustrate and analyze the required conditions for highly efficient and wavelength-tunable DW generation in PCFs between 800 – 950 nm center wavelength. The nonlinear pulse propagation in PCFs is numerically implemented using the generalized nonlinear Schrödinger equation (GNLSE)

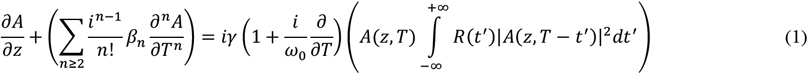

where *A*(*z, t*) denotes the complex field amplitude of the pulse and *β*_*n*_ the *n*-th order dispersion coefficients originated in the Taylor expansion of the propagation constant *β*(*ω*) at the center frequency *ω*_0_ [32]. The left-hand side of Eq. (1) accounts for dispersive effects, which are included up to the 11^th^ order in the subsequent simulations. Conversely, the right-hand side includes nonlinear effects, i.e., self-phase modulation (SPM), self-steepening (SS), and stimulated Raman scattering (SRS). Here, *γ* = *ω*_0_*n*_2_/*c*_0_*A*_*eff*_ denotes the nonlinear coefficient, defined by the nonlinear refractive index *n*_2_, the vacuum speed of light *c*_0_ and the PCFs effective mode-area *A*_*eff*_. The Raman coefficient *R*(*t*) includes SRS, taking into account both the instantaneous electronic and delayed molecular response of fused silica, implemented as outlined in Ref. [32]. The GNLSE in the form of Eq. (1) is widely used to accurately describe the nonlinear spectral broadening of ultrashort laser pulses and supercontinuum (SC) generation in PCFs as shown e.g., by Dudley *et al*. and Sylvestre *et al*. in Ref. [33,34].

To numerically illustrate the required conditions for wavelength-tunable DW generation, Fig.1 (a) and (b) show the frequency- and time-domain pulse evolution, respectively, in a 40 mm long segment of PCF (NKT, LMA-PM-5) with *A*_*eff*_ = 4.4 µ*m, γ* = 10 *W*^−1^*cm*^−1^ at 1064 nm and a zero-dispersion wavelength *λ*_*ZDW*_ close to 1060 nm. The input pulse is sech^2^-shaped and Fourier-transform limited with 100 fs pulse duration, a center wavelength of 1050 nm, a pulse energy of 12 nJ and 105 kW peak power.

**Fig. 1.**
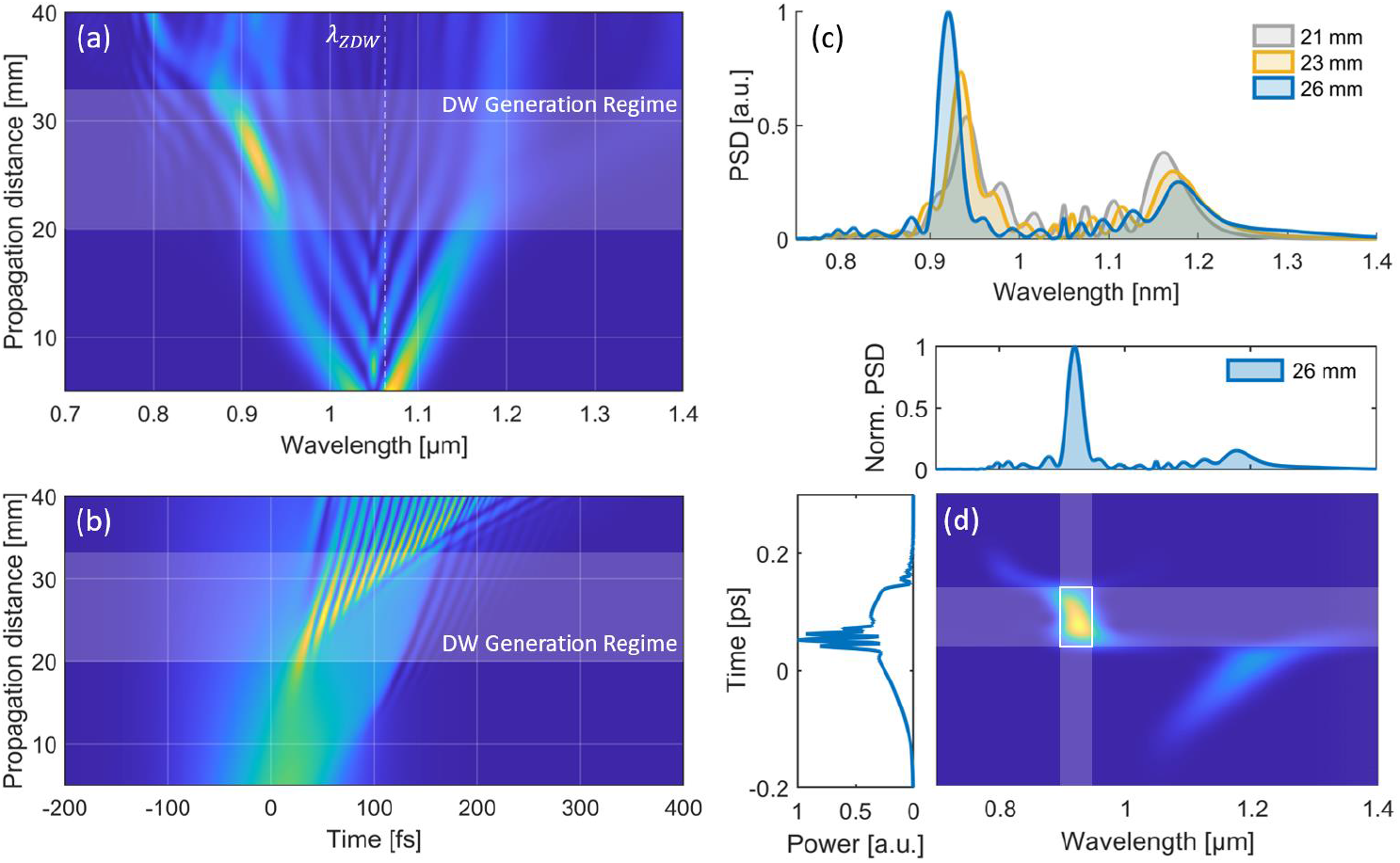
Numerical simulation of PCF dispersive wave generation. (a): Pulse evolution in the spectral domain as function of the PCF propagation distance with indicated DW generation regime. (b): Corresponding time-domain pulse evolution. (c): SC spectra at selected PCF propagation distances. (d): Spectrogram of the simulated SC at a propagation distance of 26 mm.

As shown, SPM-dominated symmetric spectral broadening is observed up to ∼20 mm of propagation. As the spectrum extends into longer wavelengths and the PCF’s anomalous dispersion regime, soliton self-compression and subsequent soliton fission occur, separating higher-order solitons into fundamental solitons. This process drives efficient energy transfer to ∼940 nm, with an emerging peak that blue-shifts toward ∼880 nm between 20 mm and 32 mm of propagation (marked in Fig. 1(a) and (b) as the DW Generation Regime).

To gain further physical insight into the origin of this energy transfer, Fig. 1(b) shows SC spectra at selected propagation distances (21–26 mm), showing that the blue-shifting peak at ∼930 nm in the normal dispersion regime of the PCF is accompanied by a reduction and red-shift of the spectral power located between 1.1 – 1.2 µm in the anomalous dispersion regime. Additionally, Fig. 1(d) depicts the spectrogram of the pulse at 26 mm propagation distance, showing that the power spectral density (PSD) peak around ∼930 nm corresponds partially to the time-domain self-compressing solitons in the pulse center and a dispersed pedestal at the pulse front. Taking these observations into account, the energy transfer to the shorter wavelengths can be explained with highly-efficient DW generation (also known as Cherenkov radiation), i.e., the resonant transfer of energy from a soliton (here at ∼1.15 µm) to a phase-matched DW in the normal dispersion regime (at ∼930 nm in this example) in the presence of higher-order dispersion [32]. Mathematical verification and further physical insight into the mechanism which ultimately enables wavelength-tunable DW generation can be obtained by analyzing the DW phase matching condition given by the relation

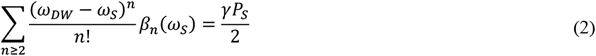

where *ω*_*S*_ and *ω*_*DW*_ represent the angular center frequencies of the soliton and DW, respectively, *β*_*n*_(*ω*_*p*_) denotes the *n*-th order dispersion coefficient at *ω*_*S*_, *γ* is the nonlinear parameter and *P*_*S*_ the soliton peak power [35]. According to Eq. (2), phase-matched DW generation at *ω*_*DW*_ occurs when the linear DW phase shift equals both linear and nonlinear phase-shift of the soliton located at *ω*_*S*_.

Using the previous numerical parameters, Fig.2 (a) shows the calculated phase-matched DW wavelength as function of the corresponding soliton wavelength for varying *P*_*S*_, obtained by solving Eq. (2). As shown, a shift of the soliton towards longer wavelengths results in a blue-shift of the phase-matched DW wavelength, in agreement with the observed numerical results in Fig.1 (c). While the decreasing *P*_*S*_ (shown e.g., in Fig.1 (c) at ∼1.15 µm) simultaneously results in a slight blue-shift, this effect on the phase-matched DW wavelength is small compared to the soliton red-shift, resulting in a net blue-shifting DW.

**Fig. 2.**
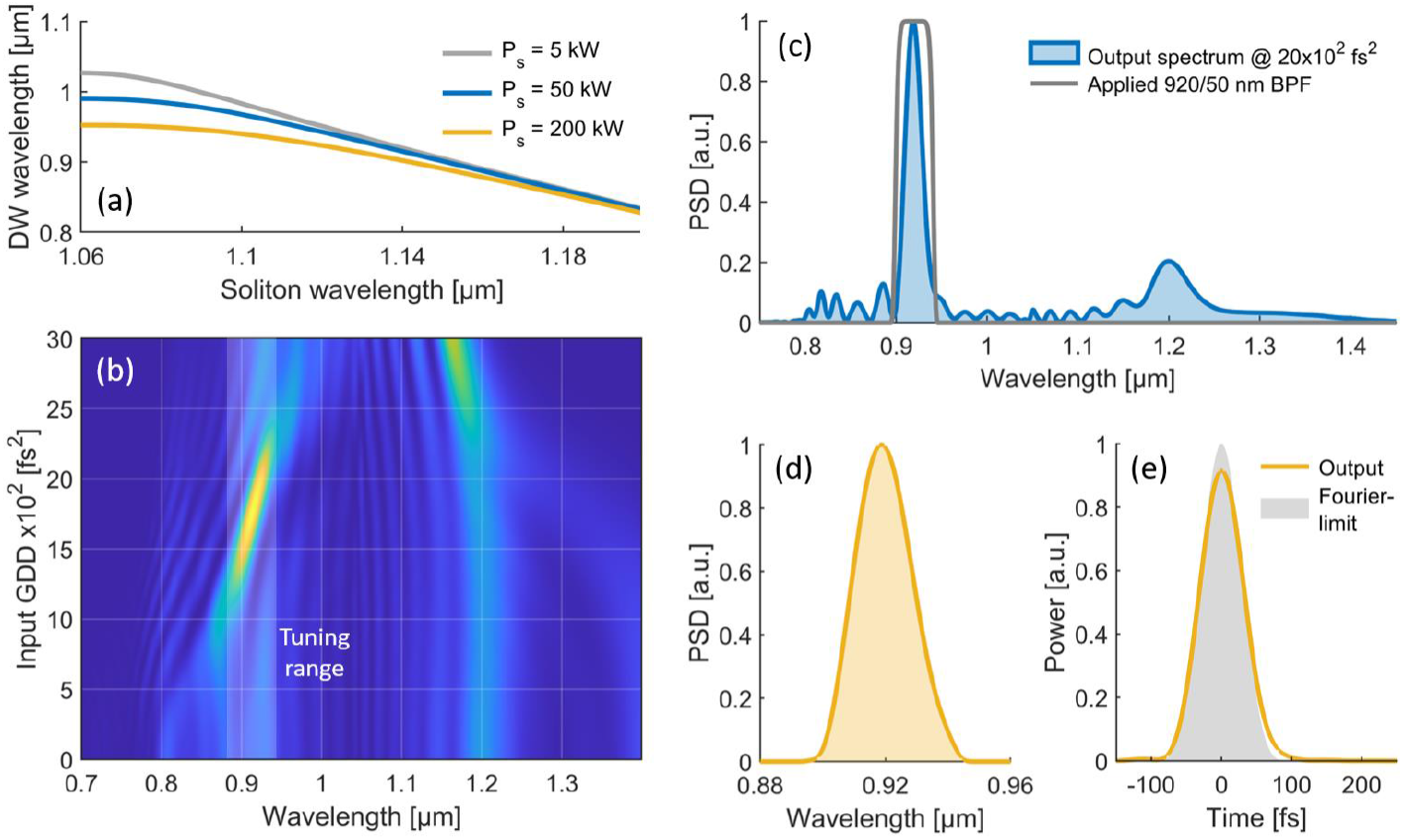
Simulation of DW phase-matching, wavelength-tunability and output pulse quality. (a): Calculated phase-matched DW wavelength as function of the soliton wavelength for varying soliton peak power. (b): Simulated PCF output spectra as function of the input pre-chirp with highlighted wavelength-tunable DW range. (c): Output SC spectrum with indicated 920/50 nm BPF to isolate the DW. (d): Spectrum of the filtered DW with a FWHM of ∼20 nm. (d): Corresponding filtered DW pulse in time-domain with a FWHM of 76 fs, compared to the Fourier-limited pulse calculated from the filtered spectrum.

The numerical analysis so far demonstrates, that wavelength-tunable DW generation can be achieved by utilizing the phase-matching condition and the resulting continuous blue-shift of the DW peak to control its center wavelength at the point of ejection from the PCF. In practice, this can be achieved by adjusting the input pre-chirp and peak power while keeping the PCF length and input power constant. To illustrate this numerically, Fig.2 (b) shows the output SC after 35 mm of PCF propagation for varying input group delay dispersion (GDD), using the same simulation parameters as before but with a slightly increased pulse energy of 13 nJ, corresponding to an initial peak power of ∼114 kW. As shown, varying the input GDD from 12.5 ∗ 10^2^ *fs*^2^ to 22.5 ∗ 10^2^ *fs*^2^ enables to shift the peak of the ejected DW peak from ∼880 nm to 940 nm in this exemplary simulation. The conversion efficiency from the 1050 nm input pulse to a 50 nm spectral band around the DW center wavelength remains larger than 40% in the complete tuning window, and reaches more than 50% around 920 nm with 6.8 nJ pulse energy. It should be noted, that for a fixed PCF length, the DW generation further depends on the input pulse energy and center wavelength. In the subsequent experimental part, this allows to optimize and fine-tune the DW conversion efficiency at the targeted center wavelength.

Besides the available pulse energy within the tuning window, a key factor for the DW output (in particular for 2PM applications) is the achievable pulse quality and duration, which directly impact the peak power delivered to the sample. Fig.2 (c) shows the output SC with ∼ 20×10^2^ fs^2^ input GDD and a DW peak centered at ∼920 nm. The indicated application of a 920/50 nm bandpass filter (BPF) results in the extraction of and ultrashort pulse shown in Fig. 2 (d) and (e) in spectral- and time-domain, respectively. With a spectral full width at half maximum (FWHM) of ∼22 nm, the DW output has a pulse duration of ∼70 fs with a clean pulse shape (i.e., no pedestals or satellites), with is close to the Fourier-limited pulse duration of 64 fs (shown in Fig.2 (e)). This can be explained with the propagation of the DW in the normal dispersion regime, which results in a subtle positive chirp that is also visible in the simulated spectrogram shown in Fig.1 (d). Similar performance is observed over the whole DW tuning range, as later also verified in the experiment. The numerically indicated parameter space of the wavelength-tunable DWs with sub-100 fs pulse duration, clean pulse quality and more than 5 nJ pulse energy seems well-suited for versatile deep-tissue 2PM applications.

## 3. Experimental Results and Discussion

### 3.1 Laser and Imaging Setup

Guided by the numerical results, an experimental setup is constructed to enable the application of wavelength-tunable DW generation for multicolor deep-tissue two-photon brain imaging. The experimental setup, shown in Fig.3 (a), consists of a laser system and a two-photon imaging capable scanning microscope. The laser system includes a home-built ultrafast Yb:fiber laser (YFL) with 30 MHz repetition rate and 2 W average power, corresponding to ∼67 nJ maximum pulse energy. The YFL output pulse train then passes through a tunable bandpass filter (TBPF, 25 nm FWHM) and a transmission grating pair (1000 lines/mm, Coherent T-1000-1040 Series), which enable adjustment of center wavelength and pre-chirp, respectively. Fig.3 (b) shows the resulting output spectrum of the YFL, together with the filtered PCF input spectrum with the TBPF in default position, centered at 1050 nm with a FWHM of ∼42 nm. Rotating the TBPF to adjust the center wavelength by ± 4 *nm* allows to fine-tune the DW generation efficiency and wavelength position for a given applied pre-chirp as mentioned before. At this state, the filtered pulses have an average power of 1.3 W with pulse energy ∼43 nJ. Fig.3 (c) shows the corresponding measured autocorrelation (AC) trace of the fully compressed pulse with a FWHM of 46 fs assuming a Gaussian pulse-shape, compared to the Fourier-transform limited pulse with 45 fs FWHM. After the GP for pre-chirp adjustment, the filtered pulse train then passes a half-waveplate (HWP) to rotate the polarization parallel to the fast axis of a 30 mm long PCF (NKT, LMA-PM-5) that is used for the DW generation. With a beam diameter of ∼1.5 mm, mode-matching with a 4 mm aspheric focus lens (FL, Thorlabs C340TME-B) enables a high PCF coupling efficiency of ∼75%. After propagation through the PCF, the beam is collimated using a 4.5 mm collimation lens (CL, Thorlabs C230TMD-B), resulting in an output beam diameter of ∼0.9 mm. A beam expander (2.7X) in front of the scanning microscope (Lens L1: 75 mm focal length, L2: 200 mm) increases the beam diameter to ∼2.4 mm. In addition, an attenuator (ATT) in form of a variable neutral density filter is implemented, allowing to dynamically adjust the average power on the sample to keep a consistent fluorescence level during deep-tissue imaging.

**Fig. 3.**
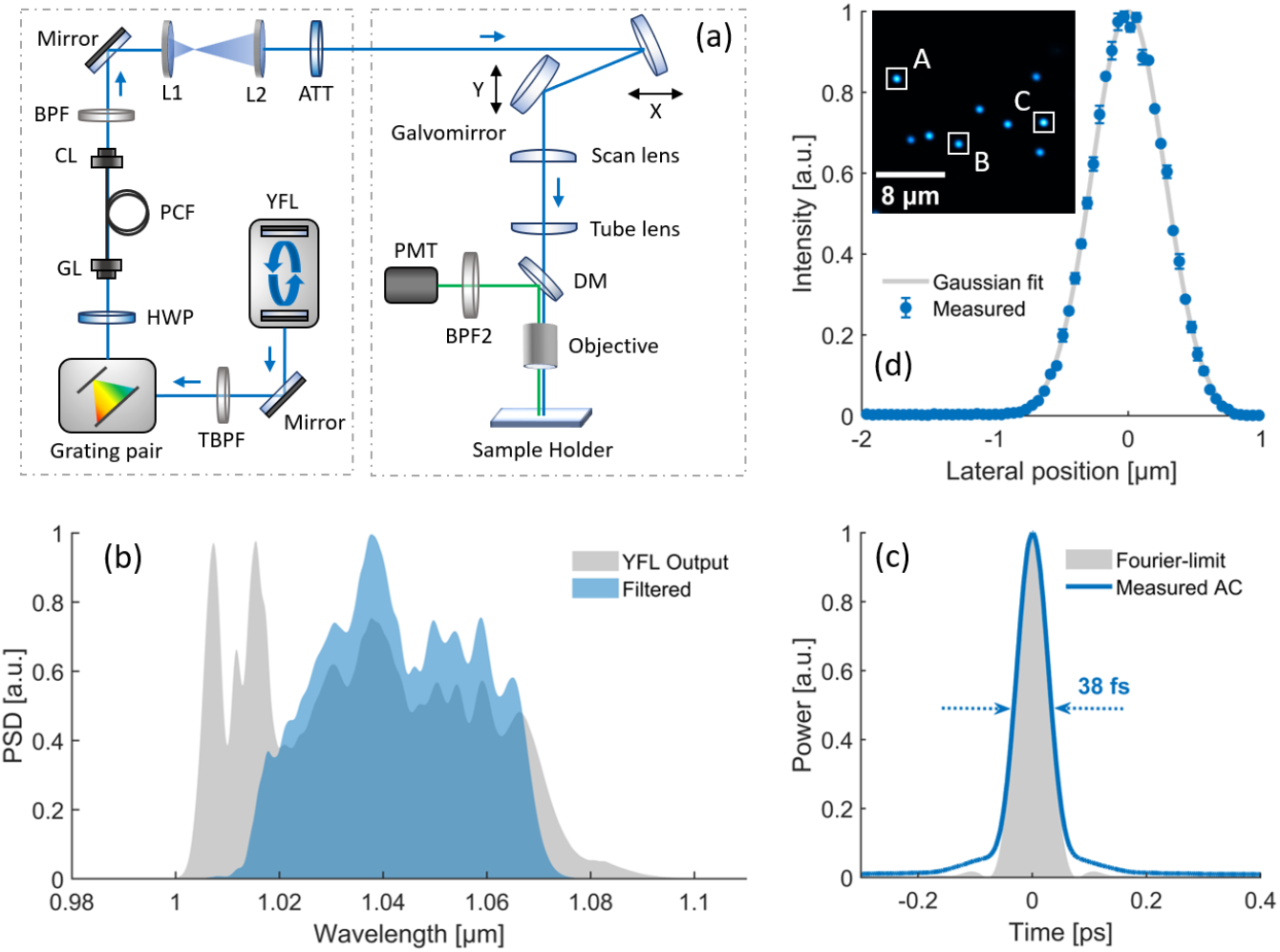
(a): Experimental setup for wavelength-tunable DW generation and two-photon neural imaging. TBPF, tunable bandpass filter; HWP, half-wave plate; FL, focus lens; CL, collimation lens; L, lens; ATT, attenuator; DM, dichroic mirror; PMT, photomultiplier-tube. (b): Output spectrum of the YFL (gray) with together with the 1050/50 nm filtered spectrum used as PCF input (blue). (c): Measured autocorrelation trace of the filtered PCF input (blue) compared to the calculated Fourier-transform-limit (gray). (d): Measured lateral PSF, determined from three sub-resolution microspheres highlighted in the inset.

The scanning microscope (Thorlabs MPM-2PKIT) used for two-photon brain imaging includes an 8 kHz resonant scanner and a ∼30 Hz mirror galvanometer to rapidly scan the field of view (FOV). The objective is a 25x water-immersion objective (Olympus XLPLN25XWMP2) with NA of 1.05, a working distance of 2 mm and a free back-aperture diameter of ∼12.5 mm. With a 5X magnification of the scan and tube lens, the input beam-diameter is therefore matched to the objectives back-aperture to ensure optimal spatial resolution. The insertion loss of the from the input of the microscope to the sample plain is ∼40%. A dichroic mirror (DM, Semrock FF665-Di02-25×36) separates the excitation beam from the epi-collected two-photon emission. An additional bandpass filter (BPF), matched to the excitation peak of the respective fluorophore, further isolates the fluorescent signal before it is detected with an amplified photomultiplier tube (PMT, Thorlabs PMT2101), which has a maximum excitation sensitivity at ∼580 nm. The sample holder is mounted on a motorized XY-stage and the z-direction is precisely adjustable using translation stage driven by a software-controlled stepper motor actuator (Thorlabs Z825B). Data and image acquisition are performed using the systems commercial software (Thorlabs ThorImageLS) and its integrated data acquisition card (National Instruments PCIe-6321).

To quantify the spatial resolution of the 2PM system, the point-spread function (PSF) is measured by imaging sub-resolution fluorescent beads (Invitrogen PS-Speck Green) with 175 nm diameter, embedded in 2% agarose with a dilution of 1:100, at an excitation wavelength of 920 nm. An acquired image of the beads is shown in the inset of Fig.3 (d), in a 23.5×23.5 µm FOV with 0.046 µm pixel size. Using the normalized fluorescent intensity profile of the three labeled beads (A, B and C), the average PSF is determined as shown in Fig. 3 (d). The calculated FWHM of the PSF (and therefore lateral resolution) allows to estimate the lateral resolution to be ±.± ± 21 *nm* (SEM), which is close to the theoretical diffraction limit of ±4± *nm*.

### 3.2. Wavelength-Tunable Dispersive Wave Generation

The described laser setup is used to generate energetic DWs at tunable center wavelengths through adjustment of the pre-chirped pulse duration, average power *P*_±*n*_ which includes the 75% coupling efficiency, and center wavelength *λ*_*c*_ of the PCF input pulse. As shown in Fig.4 (a) together with the corresponding input parameters, this approach enables SC generation in the PCF with continuously tunable DW peak in the wavelength range from 880 – 950 nm. Here, the pulse duration at the PCF input represents the FWHM of the measured AC trace, assuming a Gaussian pulse shape, and the center wavelength is determined considering the 10-dB spectral bandwidth. As shown, while the generated SCs span more than an octave for each working point, it is clearly visible that the DW generation enables highly efficient energy transfer to the phase-matched wavelength range predicted by the numerical simulations. At the PCF output, a BPF with 25 nm FWHM and fitting center wavelength is applied to spectrally isolate the DW peaks. The resulting, filtered spectra are shown in Fig. 4 (b) in steps of 10 nm, and verify the broad wavelength-tunability of the laser system within the range of 880-950 nm.

**Fig. 4.**
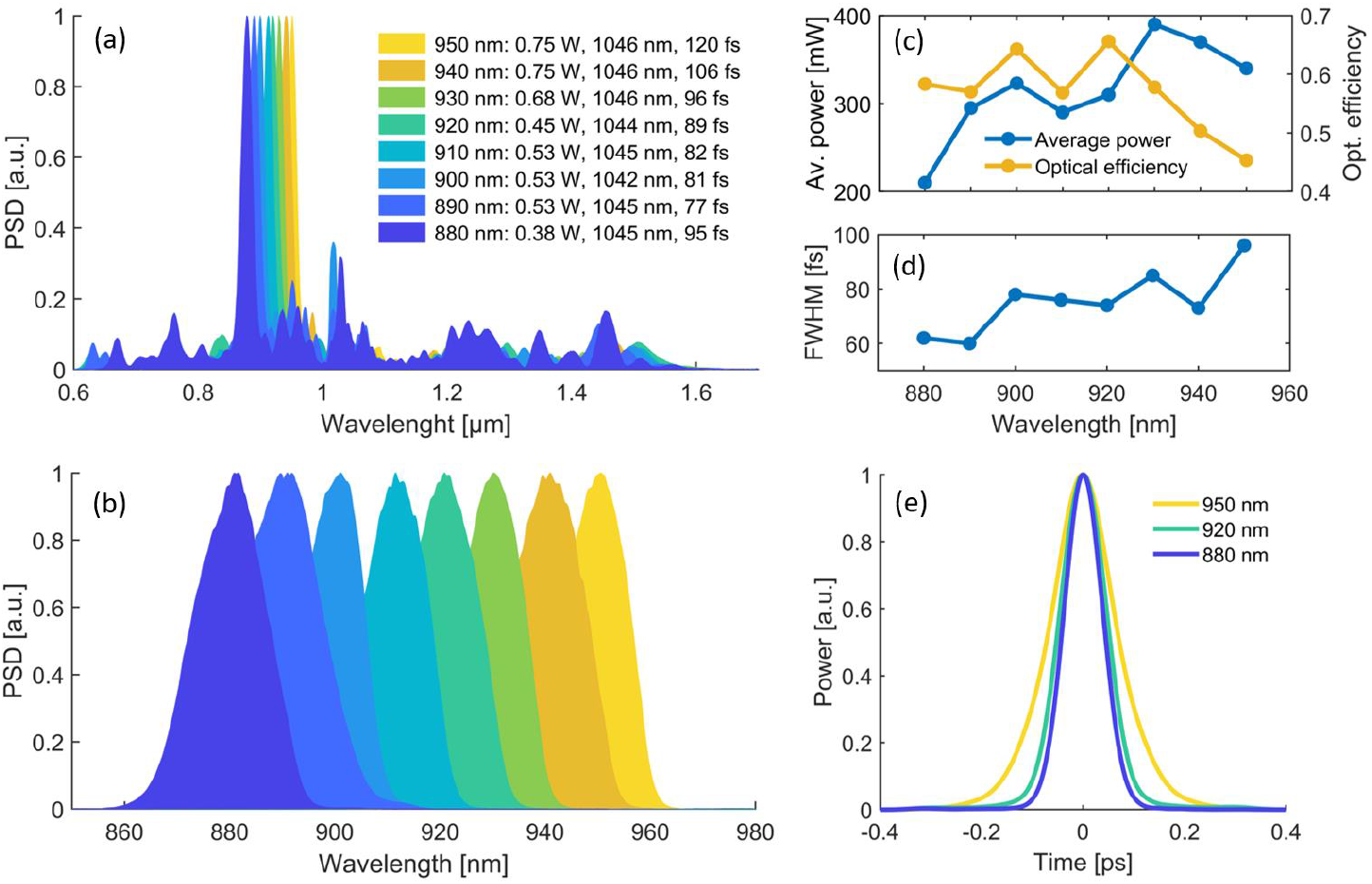
Experimental results of wavelength-tunable DW generation. (a): Measured SCs generated in the PCF with input parameters (coupled average power with 75% conversion efficiency, center wavelength, pulse FWHM assuming Gaussian pulse shape) with DW peak wavelength varying in 10 nm steps between 880 nm and 950 nm. (b): Spectra of the filtered DW peaks. (c): Corresponding measured output average power and optical conversion efficiency from the coupled PCF input power. (d): Time-domain FWHM of the corresponding measured AC traces, assuming a Gaussian pulse shape. (e): Measured AC traces of the filtered DWs at selected center wavelengths.

Fig.4 (c) shows the measured output average power and the optical conversion efficiency with respect to the average power coupled into the PCF, corresponding to the filtered DW peaks shown in Fig.4 (b). Over the complete tuning range (880-950 nm), pulses are generated with more than 200 mW average power ∼6.7 nJ pulse energy. At 930 nm center wavelength, the average power reaches a maximum of up to 380 mW with 12.7 nJ pulse energy. The optical efficiency is simultaneously larger than 45% over the full tuning-range, reaching up to 66% at 920 nm.

As further shown in Fig.4 (d), the pulse duration of the filtered DW peaks (determined as the temporal FWHM of the measured AC traces, assuming a Gaussian pulse shape) remains below 100 fs over the complete tuning range, reaching down to 60 fs at 890 nm center wavelength. Fig.4 (e) shows the measured AC traces for selected DW center wavelengths, verifying a clean pulse shape without pedestals or satellite pulses. The demonstrated parameter space of the system, i.e., sub-100 fs pulses at more than 6.7 nJ pulse energy between 880-950 nm, is well-suited for deep-tissue neural two-photon imaging applications.

### 3.3. Deep-Tissue Two-Photon Imaging at 920 nm in EGFP-labeled Mouse Hippocampus

As a first step to demonstrate flexible deep-tissue two-photon imaging capability, the wavelength-tunable DW generator is applied to visualize neuronal and vascular structures in the hippocampus of a transgenic mouse model that ubiquitously expresses EGFP in all the cells with an excitation peak at ∼920 nm [36]. To match the excitation peak of EGFP, the DW generator is tuned to 920 nm center wavelength (adjusted for maximum excitation while avoiding photobleaching and photodamage), an average power of 310 mW (∼10 nJ pulse energy) and ∼74 fs pulse duration, as shown in Fig.4. A 520/36 nm BPF (EO #67-030) is used in front of the PMT (Fig.3 (a)) to isolate the EGFP emission.

Fig. 5 (a) shows a depth scan in the mouse hippocampus to an imaging depth of up to 600 µm (in steps of 100 µm). Imaging is performed in a 443.3×443.3 µm FOV (1024×1024 pixels), a corresponding pixel size of 0.433 µm and a pixel dwell time of 0.122 µs. To reduce background noise, each image represents the average of 10 consecutive frames, taken at a frame-rate of ∼10 frames/second with an overall acquisition time of ∼5 seconds (including the processing time of the system). To keep a constant two-photon fluorescence level for increasing imaging depth *z*, the average power on the sample is continuously increased during the scan using the ATT, from an initial 8 mW at *z* = 0 µ*m* (28 mW at the system input) to 59 mW at the maximum depth *z* = 600 µ*m* (∼168 mW at the system input).

**Fig. 5.**
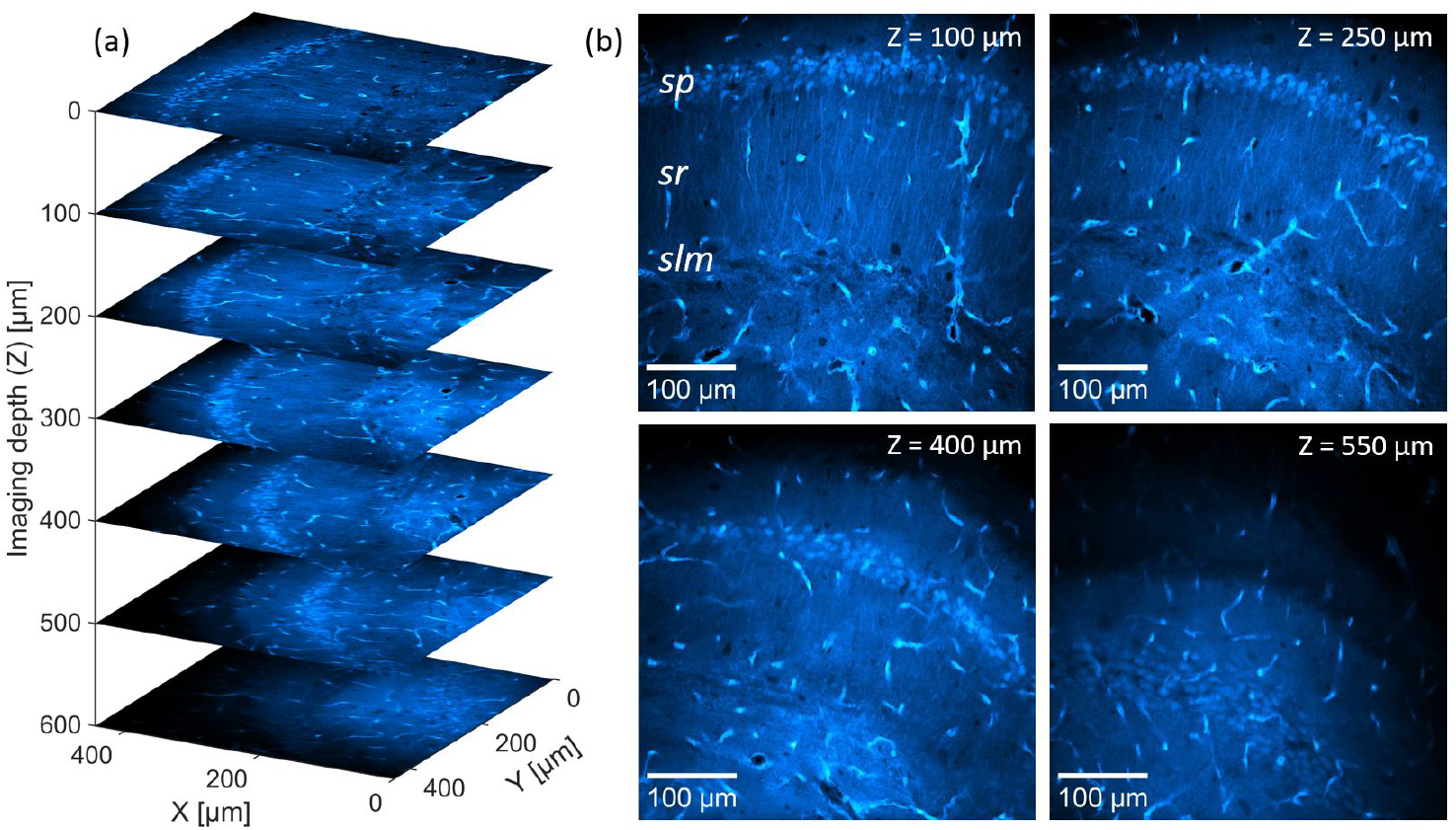
Deep-tissue imaging in EGFP labeled mouse hippocampus, using the DW generator at 920 nm center wavelength. (a): Depth scan of the EGFP-labeled brain slice (concretely the CA1-hippocampal region), in 100 µm steps to a maximum depth of 600 µm, in a FOV of 443×443 µm (1024×1024 pixels). The soma of the pyramidal neurons (in the *sp*) and their projections (in the *sr* and *slm*) are clearly observed alongside some vascular structures. (b): Corresponding images at selected depths, shown in the same FOV. CA1, *cornu ammonis 1; sp, stratum pyramidale; sr, stratum radiatum; slm, stratum lacunosum-moleculare*;

To further illustrate the resolved neuronal structure at increasing depth, Fig.5 (b) shows obtained images from the depth scan at selected imaging depth. Up to *z* = 250 µ*m*, distinct layers of pyramidal neurons of the CA1 region with well-defined soma and their projections in the *sr* and *slm* are clearly visible, along with selected blood vessels. While the cell bodies remain distinguishable at greater depths, neuronal projections become progressively less visible (as shown e.g., for z = 400 µm in Fig.5 (b)). Representing a characteristic feature of 2PM, this observation can be explained by the increased background noise and image blurring, which arise from enhanced two-photon emission at the sample surface due to the larger average power at increasing imaging depths [37,38].

### 3.4. Deep-Tissue Two-Photon Imaging at 950 nm in SYTOX Orange labeled Mouse Cerebellum

To demonstrate wavelength-tunable deep-tissue 2PM capability, the DW generator is further applied to visualize and distinguish individual neuronal nuclei in a stained mouse brain cerebellum. In this case, the used brain sample is a prepared slide (SunJin Lab FluoTissue) containing a 550 µm thick mouse brain section, multi-labeled with Alexa Fluor 488 for the blood vessels, SYTOX Orange for the neuronal nuclei and Alexa Fluor 647 for dopaminergic neurons. The sample is mounted with an optical clearing reagent (SunJin Lab RapiClear 1.52) and sealed between a microscopy slide and a glass cover slide. Two-photon imaging is performed with the DW generator tuned to 950 nm center wavelength for efficient excitation and visualization of the SYTOX Orange labeled neuronal nuclei [39]. The maximum available average power at this working point is 340 mW (∼11 nJ pulse energy) and the pulse duration is ∼96 fs, as shown in Fig.4 (c) and (d), respectively. To isolate the emission spectrum of SYTOX Orange, a 572/28 nm BPF (EO #84-100) is installed in front of the PMT as indicated in Fig.3 (a).

To verify the ability for deep-tissue imaging using the 950 nm DW generator output, Fig.5 (a) depicts the result of a depth scan of the mouse brain slice in the cerebellum area down to a maximum imaging depth of 450 µm in 100 µm steps. Each individual image is acquired in a 595×594 µm FOV (1024×1024 pixels) with ∼0.581 µm pixel size and a pixel dwell time of ∼0.2 µs as the average of 10 consecutive frames (∼10 frames/s with ∼5 seconds acquisition time). To keep the fluorescence level on the PMT similar for increasing depth, the average power is again adjusted via the ATT, from ∼23 mW (57 mW at the input) at *z* = 0 µ*m* to 76 mW (190 mW at the input) at *z* = 450 µ*m*. As shown, the obtained depth scan allows to clearly resolve neuronal nuclei and scattered blood vessels in the molecular and granular layer of the mouse cerebellum, down to the full imaging depth of 450 µm, which corresponds to the overall sample thickness.

The nuclei of cerebellar granule cells in the granular layer of typical mouse models have a diameter of ∼5 µm (well above the measured lateral resolution of the 2PM system) and almost entirely occupy the neuronal cell bodies [40]. To characterize the ability of the system to resolve the individual nuclei at increasing imaging depths, Fig.6 (b) shows imaged sections obtained at *z*=50 μ*m* and *z*=450 μ*m* with marked regions of interest (ROI, labeled as A and B). The ROIs A and B are depicted separately with 2.9X magnification, corresponding to a 208×208 µm FOV (358×358 pixels). ROI A, located at a shallow depth of *z* = 50 µ*m*, shows clearly resolved nuclei of individual neurons in the granular layer. In comparison, a reduction in contrast due to the increased background noise can be observed in ROI B, imaged at the maximum depth of *z* = 450 µ*m*.

**Fig. 6.**
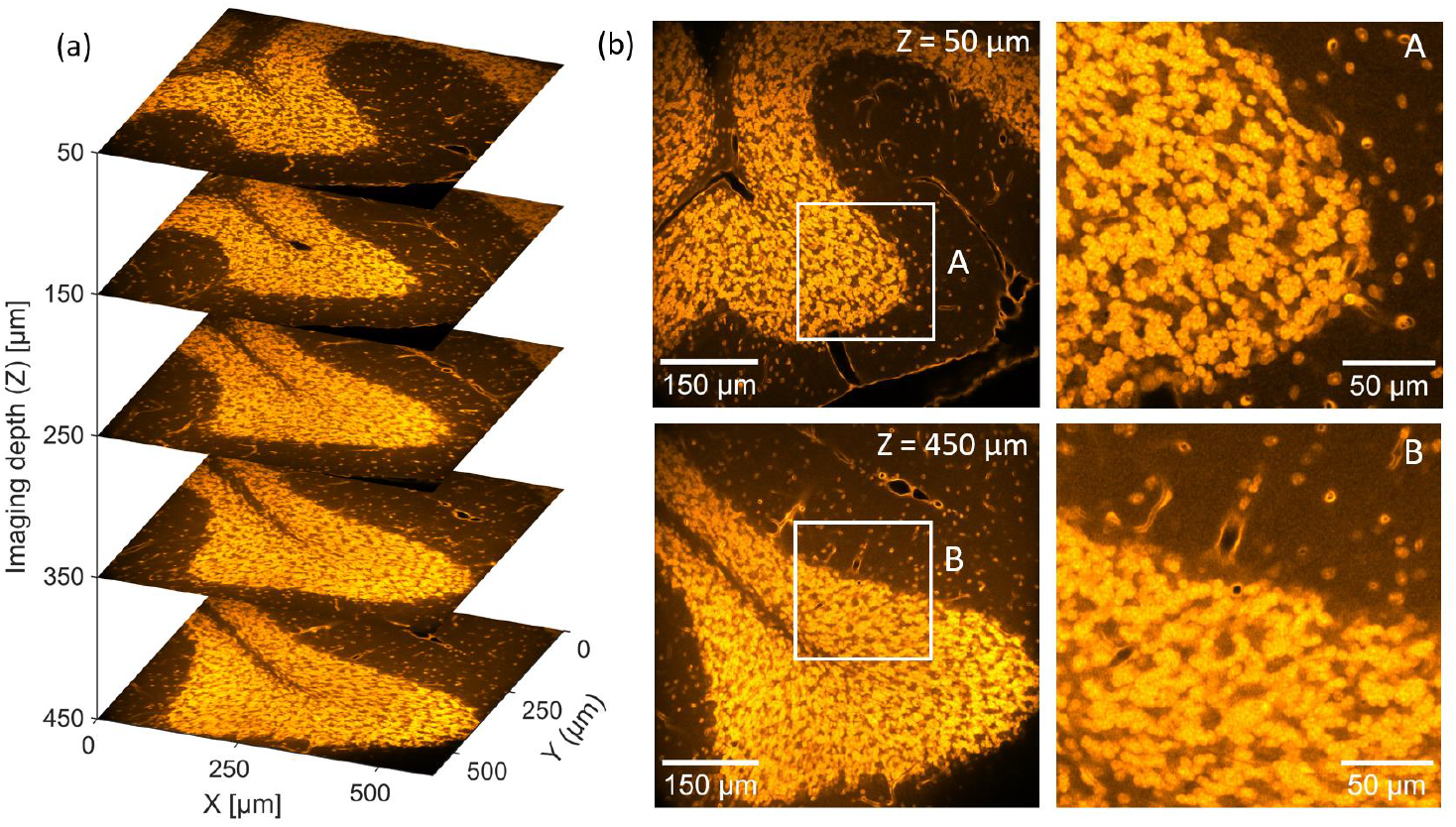
Deep-tissue imaging in SYTOX Orange labeled mouse cerebellum using the 950 nm output of the DW generator. (a): Depth scan of the SYTOX Orange labeled brain slice (showing an area of mouse cerebellum) taken in 100 µm steps down to a maximum depth of 450 µm, covering a 595×595 µm FOV (1024×1024 pixels) (b): Images obtained at 50 µm and 450 µm depth in a 595×595 µm FOV (1024×1024 pixels), with marked and highlighted ROIs (A, B) magnified with a zoom factor of ∼2.9X to a FOV of 208×208 µm (358×358 pixels), to visualize the ability to resolve individual granule cell nuclei for increasing depth.

Nevertheless, the granular nuclei remain distinguishable even at this depth, demonstrating effective imaging performance throughout the entire depth range. This experiment therefore clearly showcases, that the 950 nm DW driven imaging setup enables sufficient resolution and contrast for high-performance deep-tissue imaging of fine neuronal structures within intact brain tissue. Notably, compared to the images of the EGFP-labeled mouse brain in Fig. 5, the SYTOX Orange-labeled sample exhibits visibly improved contrast and image quality. This enhancement can be attributed to additional tissue clearing, as well as the highly specific labeling of neuronal nuclei by SYTOX Orange compared to the ubiquitously expressed EGFP, resulting an improved signal-to-background ratio.

## 5. Conclusion

This work presents a versatile, fiber-optic, wavelength-tunable laser system that delivers high-energy ultrashort pulses tailored for versatile deep-tissue two-photon brain imaging. By utilizing precisely controlled dispersive wave generation in photonic crystal fibers, the constructed achieves sub-100 fs pulse durations with pulse energies exceeding 6.7 nJ across an 880–950 nm spectral range, with record optical conversion efficiencies of up to 65% from an Yb:fiber laser. Comprehensive numerical simulations are applied to verify and investigate the underlaying physical mechanism. Our approach enables excitation-matched two-photon imaging for various fluorophores and neuronal markers, achieving high-resolution visualization of neuronal structures and vascular networks at significant imaging depths. Two-photon imaging at 920 nm in EGFP-labeled hippocampal tissue enables detailed visualization of neuronal morphology to depths of 600 µm, while imaging at 950 nm allows to resolve individual neuronal nuclei in SYTOX Orange labeled cerebellar tissue up to 450 µm depth. These imaging results highlight the system’s capability to dynamically match the excitation profiles of different fluorophores to ensure optimal image contrast, signal-to-noise ratio, and minimized photodamage. The demonstrated combination of broad tunability, compact design, and high optical efficiency further establishes this fiber-optic laser platform as a highly adaptable tool for advanced neuroimaging, offering significant potential for exploring neural connectivity, synaptic dynamics, and brain activity in both healthy and pathological conditions

## A. Methods

### Animals

The mouse model is an adult wild-type C57BL/6J male mouse (∼6 months old, P180–P210) with ubiquitous EGFP expression. No experiments with living animals were performed. The animal breeding and euthanasia to obtain the tissue were approved by the ethical research committees of respective national/local authorities: *Freie und Hansestadt Hamburg, Behörde für Gesundheit und Verbraucherschutz*, Hamburg, Germany (ORG1108). Sampling was performed following the protocols and guidelines of the Ethical Committee and the directives of the European Union Council 86/609 and 2010/63.

### Sample Preparation

The brain of the EGFP mouse model was quickly extracted from the skull and fixed with 4% paraformaldehyde for 24h in the fridge. The following day, the tissue was washed with phosphate buffer 1M three times. Subsequently, the brain was sliced coronally (∼2 mm thick) with a scalpel, and then, under the microscope the hippocampus area was separated and mounted in a compartment of a glass-bottom µ-slide microscopy chamber (Ibidi, #80427-90) using Fluoromount (Invitrogen, #00-4958-02). Imaging was performed through the ∼0.5 mm thin undersurface of the slide, to enable the insertion of a purified water droplet between the sample and the water immersion objective.

## B. Funding

This work has been supported by Deutsches Elektronen-Synchrotron DESY, a member of the Helmholtz Association (HGF), POF IV DMC and the Cluster of Excellence ‘Advanced Imaging of Matter’ of the Deutsche Forschungsgemeinschaft (DFG) – EXC 2056 – project ID 390715994. A.M.-A.’s contributions were supported by the European Union’s Horizon 2020 research and innovation program through the Marie Skłodowska-Curie grant agreement No. 101030402. M.S.’s contribution was supported by the PIER Hamburg/Boston seed grant (PHM-2019-03) and PIER seed grant (PIF 2020-10) awarded by the University of Hamburg, Germany; and the Forschungsförderungsfonds der Medizinischen Fakultät grant (NWF-20/10) awarded by the University Medical Center Hamburg-Eppendorf (UKE), Germany. Additionally, M.G. would like to express their gratitude for the financial support received from the Joachim Herz Stiftung in Hamburg, Germany.

## C. Acknowledgements

We like to thank Prof. Dr. Udo Bartsch (Institute of Ophthalmology, University Medical Center Hamburg-Eppendorf, Hamburg, Germany) for kindly providing the EGFP-expressing mouse model. We are also grateful to Edda Thies (Institute of Neuropathology, University Medical Center Hamburg-Eppendorf, Hamburg, Germany) for her assistance with sample preparation and staining. Additionally, we appreciate the support of Susanna Gevorgyan and the group of Prof. Dr. Christian Betzel (Institute for Biochemistry and Molecular Biology, Universität Hamburg, Hamburg, Germany) in preparing the microsphere sample. Finally, we thank Dr. Yi Hua and Dr. Neetesh Singh (Deutsches Elektronen-Synchrotron DESY, Hamburg, Germany) for their valuable discussions and insights.

## D. Competing Interest

The authors declare no competing interests.

## E. Data availability

Data underlaying the results presented in this paper are not publicly available at this time but may be obtained by the authors upon reasonable request.

## F. Author Contributions

M.E., A.M.-A. and M.S. conceived and conceptualized the experiments. M.E. conceived and constructed the laser system and performed the laser characterization and imaging experiments. A.M.-A. and M.S. prepared the EGFP mouse brain samples. F.X.K., M.G. and M.P. supervised the project. M.E. wrote the manuscript with contributions from all coauthors.

## Notes

### Competing Interest Statement

The authors have declared no competing interest.

